# Fluid shear stress on astrocytes and microglia promotes neuronal toxicity *in vitro* via purinergic signaling

**DOI:** 10.1101/2025.04.29.651240

**Authors:** Sarah Tenney, Christopher Streilein, Andrew Bermudéz, Zackary Keller, R Chase Cornelison

## Abstract

Interstitial fluid flow plays a critical role in maintaining function and homeostasis in neural tissue, and dysregulation of this flow due to injury or disease results in mechanical stress that is associated with several neuropathologies, including traumatic brain injury, ischemic stroke, and glioma. Glial cells such as astrocytes and microglia are known to respond to mechanical forces like fluid shear stress but the impact of this stress on their functionality and any subsequent impact on neurons remains poorly defined. To investigate how pathologically high fluid shear stress modulates astrocyte and microglia function and to determine whether glial responses to fluid shear influence neuronal survival and morphology, we applied low pathological levels of fluid shear stress (0.1 dynes/cm^2^) to cultured human astrocytes and microglia and assessed functional changes including metabolic activity, metabolite release, lipid droplet accumulation, and phagocytic activity. Conditioned media from these glia were then applied to differentiated SH-SY5Y neurons to evaluate effects on cell survival and neurite outgrowth. We then focused on identifying the soluble factor mediating the observed neurotoxicity. We found that fluid shear stress promotes distinct functional responses in astrocytes and microglia, including increased metabolic activity in astrocytes, increased lipid droplet accumulation in microglia, and heightened release of extracellular ATP in both cell types. Exposure to shear-conditioned glial media significantly reduced neuronal survival and neurite length. This neurotoxic effect was abolished by activated charcoal filtration but not by boiling and was prevented through P2×7 receptor inhibition in neurons, suggesting extracellular ATP as a causative agent. These findings indicate that high fluid shear stress promotes glial-mediated neurotoxicity via purinergic signaling. This study helps to characterize glial-neuronal mechanobiological interaction in the context of neuropathology and provides support for targeting purinergic signaling pathways as a therapeutic approach for neuropathologies associated with altered interstitial fluid flow.

**Statement of significance:** Glial cells, until recently merely considered to support neurons, are now known to play critical roles in neural tissue development and function. Astrocytes and microglia play diverse roles in the central nervous system, adopting phenotypes that both promote and resolve pathology. Within brain tissue, disruptions to interstitial fluid flow are associated with brain injury, ischemic stroke, and glioma. This study identifies fluid shear stress as a modulator of glial cell function with downstream neurotoxic consequences. Namely, we found that increased release of extracellular ATP by glial cells under high shear promotes neurotoxicity via P2×7 receptor signaling. These findings help to characterize mechanobiological glial-neuronal communication and lend support for the therapeutic efficacy of P2×7 receptor inhibition in the treatment of neuropathology.

## Introduction

Glia are a general class of cells in the nervous system that play critical roles in all aspects of neural tissue development and function (1,2). The primary glial cells of the central nervous system (CNS) include astrocytes, microglia, and myelinating oligodendrocytes. Astrocytes are the most abundant cells of the central nervous system, with varied roles in ionic and metabolite regulation, extracellular matrix production, neurotrophic and synaptic support, blood-brain barrier regulation, and innate immunity (3,4). Microglia are the resident immune cell of the brain and spinal cord, primarily providing immune surveillance and effector functions but also influencing synaptic formation and pruning, neurogenesis, and remodeling of neural circuits (5,6). These cells exhibit a great deal of diversity, with phenotypic expression varying by anatomical location, disease status, and age (6–8).

Astrocytes and microglia reside within the interstitial space of the brain parenchyma, which is highly hydrated at ∼80% water. The brain therefore heavily relies on the constant perfusion or flow of interstitial fluid to maintain tissue function (9). While it is somewhat controversial whether convective flow or simple molecular diffusion occurs in the brain under normal conditions, recent research validates the possible existence of fluid convection in the brain by computational and experimental methods, which could be exacerbated by pathology (10–12). Interstitial flow is a biophysical process that replenishes nutrients and removes cellular waste. Fluid flow also exerts mechanical shear stresses at the cell surface to which cells can sense and respond through mechanosensitive receptors and channels in the cell membrane (13,14). Under physiological conditions, fluid shear stresses in the CNS are estimated on the order of 0.01 dyne/cm^2^ (15,16). Conditions such as ischemic stroke, glioma, and traumatic injury disrupt fluid flow pathways around and within the CNS, which can cause predicted increases in fluid shear stresses up to 2 orders of magnitude (17–19). These ‘pathological’ levels of fluid flow have been shown to promote inflammatory phenotypes in several cell types, including both astrocytes and microglia (20–24). Both astrocytes and microglia can adopt reactive functions ranging from neurotoxic to neuroprotective. We recently showed that direct application of fluid shear onto neurons in culture decreases cell survival and neurite length (18), but it is currently unknown if shear-stimulation of glial cells can also indirectly stimulate neurotoxic outcomes. Understanding how fluid shear stress influences glial cell function and their effects on neurons may inform therapeutic development to mitigate neurodegeneration and promote repair.

In this study we investigated the effects of high fluid shear stress on astrocyte and microglia functionality and the downstream consequences on neurons *in vitro*. We examined how fluid shear stress influences key functions of glial cells, including metabolic activity, metabolite release, phagocytic activity, and lipid droplet accumulation. We also quantified the effects of glial shear-conditioned media on neuronal cell survival and neurite extension. We identify a mechanism involving purinergic signaling through which high fluid shear on glial cells promotes neuronal toxicity *in vitro*. This effect was observed for both astrocytes and microglia despite both cell types exhibiting differential functional cellular responses to high shear by other metrics. Our results provide new insight into the potential role of fluid forces in neurodegeneration after injury or in disease and may help further explain published results showing therapeutic benefits of inhibiting purinergic signaling in myriad neuropathologies (25–31).

## Results

### High fluid shear stress alters metabolite release from glial cells

We and others have previously estimated that interstitial fluid shear stresses can range from 0.1-1 dynes/cm^2^ in inflammatory conditions like neural injury and cancer (18,23,24,32). To investigate how this level of fluid shear forces impacts glial cell function, we modeled fluid shear stress on 2D cultures of glial cells using an orbital shaker (**Figure 1A**). This model was previously validated *in silico* and *in vitro* via optical Doppler velocimetry, with a rotational speed of 100 revolutions per minute (rpm) at a defined radius of rotation equating to a low pathological shear stress of 0.1 dynes/cm^2^ (32). Using this model, we exposed astrocytes and microglia to high fluid shear stress and assessed metabolic activity and metabolite release within the conditioned media. We find that pathological shear stress induces a significant increase in metabolic activity in astrocytes, as measured by an alamarBlue assay but metabolic activity of microglia was not affected (**Figure 1B**). Additionally, astrocytes release significantly more glutamate under high shear compared to static conditions, whereas microglial glutamate release is not significantly impacted by exposure to high fluid shear stress (**Figure 1C**). The release of lactate is also not affected for either glial cell type (**Figure 1D**), but both astrocytes and microglia increase the release of extracellular adenosine triphosphate (ATP) under high fluid shear stress compared to cells in static conditions (**Figure 1E**).

**Figure 1.**
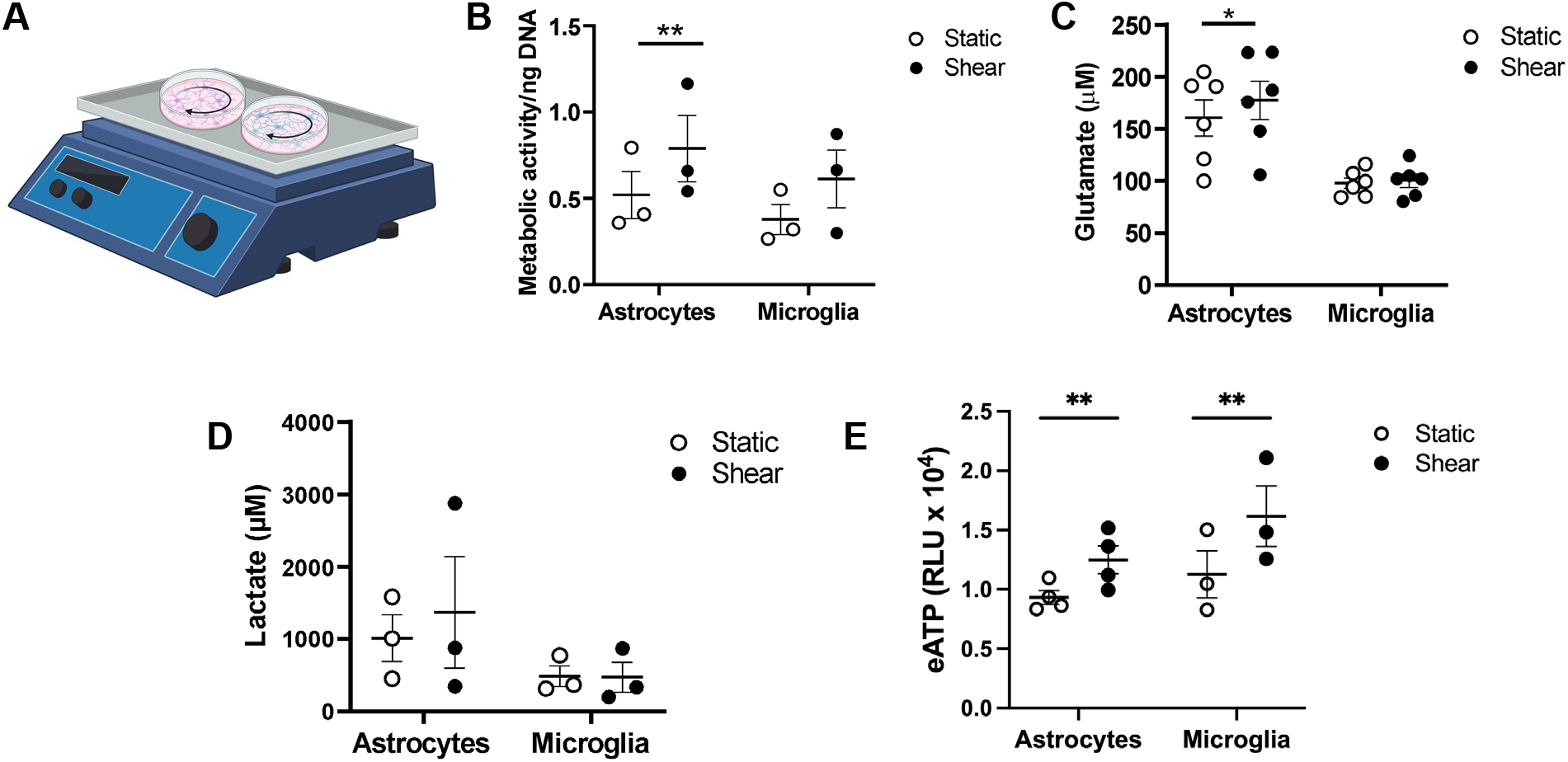
High fluid shear stress alters metabolite release from glial cells. A) Schematic of using an orbital shaker to generate 0.1 dynes/cm2 of fluid shear stress for overnight shear-conditioning of glial cells. Graphic made with an institutional license of Biorender. B) Glial cell metabolic activity after culture under static versus shear conditions by alamarBlue HTS assay. C) Quantification of glutamate release under static versus shear. D) Quantification of lactate release under static versus shear. E) Quantification of astrocyte and microglia release of extracellular ATP (eATP), presented as relative luminescence units (RLU), under static versus shear conditions. Groups were compared by ratio paired t-tests for each glial cell type. *p<0.05, **p<0.01.

### High shear enhances microglial accumulation of lipid droplets and phagocytosis

Given the established connections between aberrant accumulation of lipid droplets in glial cells and neuropathology (33–36), we used a BODIPY dye to stain lipid droplets in glial cells after one hour of exposure to 0.1 dynes/cm^2^ in a microfluidic device (33–36). Astrocytes do not increase their sequestration of lipid droplets in response to high fluid shear stress compared to cells kept under static conditions (**Figure 2A, B**). In contrast, microglia do significantly increase lipid droplet accumulation in response to shear stress (**Figure 2C, D**). We also examined particle uptake in microglia, since this is a key function of immune cells and fluid shear stress has been previously shown to increase phagocytic activity in astrocytes (37). We treated microglia from either static or sheared cultures with fluorescent 1 μm particles for 1 hour, then quantified the number of cells with particles. Like astrocytes, exposing microglia to high fluid shear stress heightens phagocytic activity compared to cells held static (**Figure 2E, F**).

**Figure 2.**
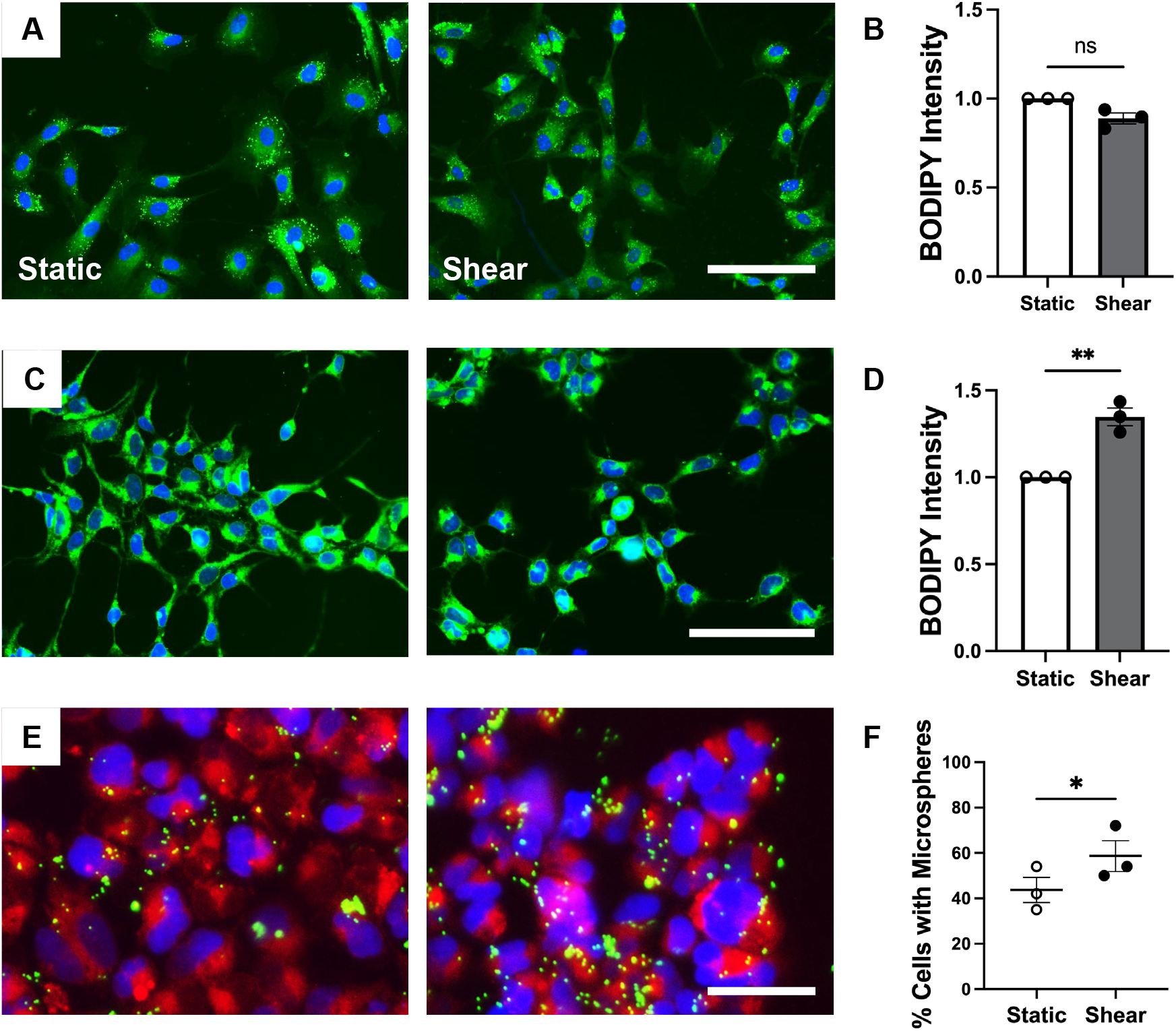
High shear enhances microglial accumulation of lipid droplets and microglial phagocytosis. A) Representative fluorescence images of BODIPY staining in astrocytes under static (left) or 0.1 dynes/cm^2^ shear (right). Scale bar is 150 µm. Green represents BODIPY dye, blue represents DAPI-stained nuclei. B) Quantification of lipid droplet accumulation in astrocytes. Normalized to static condition. C) Representative images of BODIPY staining in microglia under static (left) or 0.1 dynes/cm^2^ shear (right). Scale bar is 100 µm. Green represents BODIPY dye, blue represents DAPI-stained nuclei. D) Quantification of lipid droplet accumulation in microglia. Normalized to static condition. (E) Representative images of bead phagocytosis assay with microglia cultured under static (left) or shear conditions (right). Scale bar is 50 µm. Red represents CellTracker, green represents fluorescent microspheres, blue represents DAPI-stained nuclei. F) Quantification of microglial phagocytic activity. Groups for static and shear were compared by ratio paired t-tests. *p<0.05, **p<0.01.

### Glial shear-conditioned media reduces neuronal cell count and neurite length

We previously found that exposing neurons directly to 0.1 dynes/cm^2^ of fluid shear stress results in shorter neurite outgrowth compared to neurons held static (18). To investigate if there are additional indirect effects of fluid shear stress on neurons, we treated SH-SY5Y differentiated neurons with conditioned media from either astrocytes or microglia exposed to 0.1 dynes/cm^2^ fluid shear stress overnight. After 3 days, neurons were fixed and stained for beta-III tubulin and evaluated for cell survival and neurite length (**Figure 3A**). Representative images of treated and stained neuron cultures are shown in **Figure 3B**. Treating neuron cultures with shear-conditioned media results in significantly fewer neurons surviving compared to cells treated with conditioned media from glia held in static (**Figure 3C**). This effect is consistent for shear-conditioned media from astrocytes, microglia, and co-cultures of both astrocytes and microglia. In addition, shear-conditioned media resulted in shorter neurites compared to static glial conditioned media (**Figure 3D**). The static glial condition promotes similar neurite length to non-conditioned media controls (not shown), so we conclude shear-conditioned glial media at least prevents neurite outgrowth during the 3-day incubation period and may promote neurite retraction.

**Figure 3.**
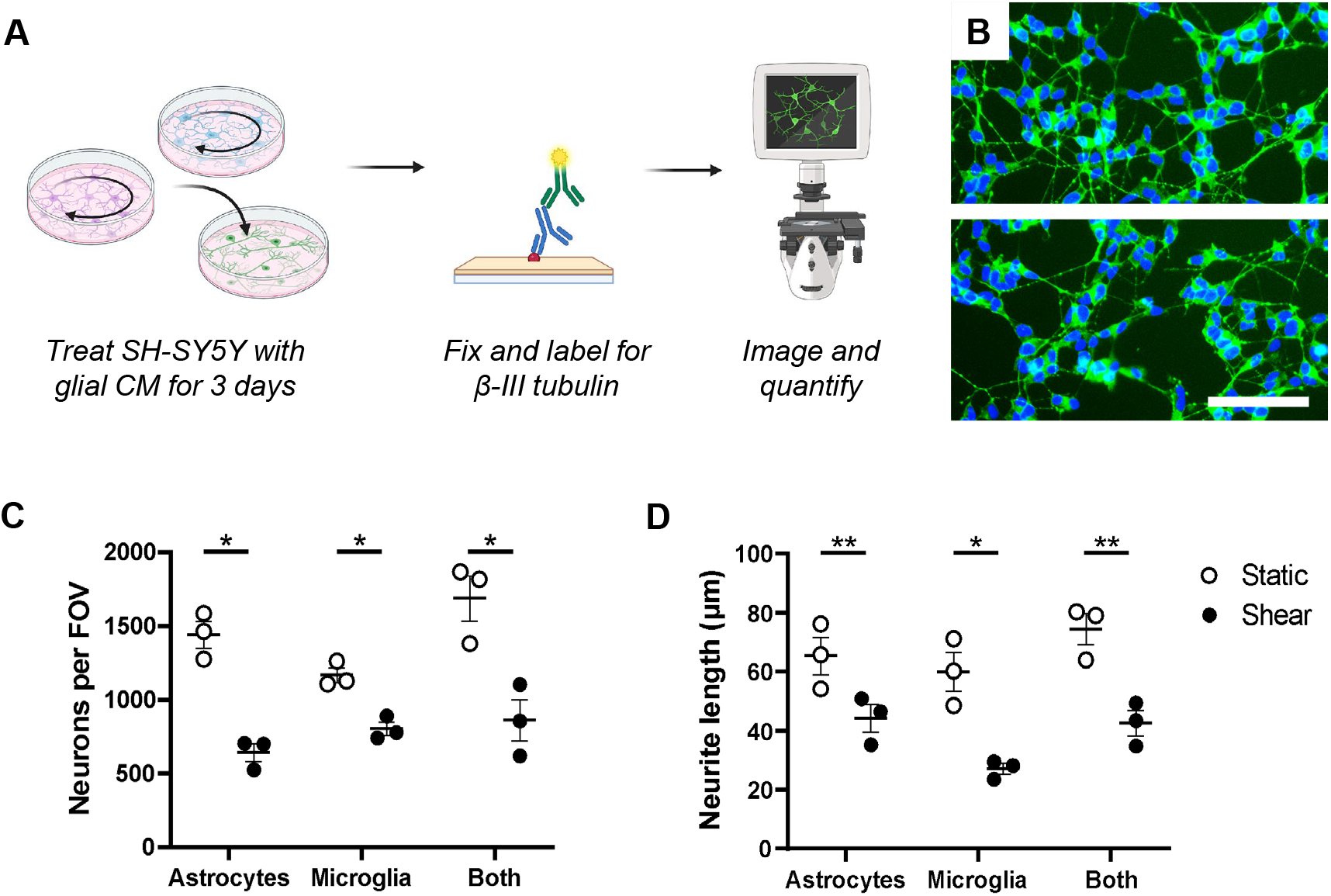
Glial shear-conditioned media reduces neuronal cell count and neurite length. A) Schematic of experimental setup, including treating differentiated SH-SY5Y neuron-like cells with glial shear-conditioned media (CM) for 3 days, staining, imaging and quantifying. Graphic made with an institutional license of Biorender. B) Representative fluorescence images of SH-SY5Y cells treated with conditioned media from microglia cultured in static conditions (top) or exposed to 0.1 dynes/cm^2^ of fluid shear stress (bottom). Green represents β-III tubulin, and blue represents DAPI-stained nuclei. Scale bar is 100 µm. C) Quantification of neuron-like cells per field of view (FOV). D) Quantification of neurite length in microns. Groups for static and shear were compared by ratio paired t-tests.

### The toxic effect of glia under high shear is removed by charcoal stripping but not boiling

Towards understanding how glial cells mediate neurotoxicity under shear, we began by boiling co-cultured glial conditioned media to denature any proteins present. If the responsible molecule is a protein, this boiling step would eliminate the neurotoxic effect. Nonetheless, boiling the conditioned media does not eliminate the effect of shear-conditioned media on neuronal cell count (**Figure 4A**) nor neurite length (**Figure 4B**). These results suggest the soluble factor responsible for neuronal toxicity is not proteinaceous. We next filtered the glial conditioned media through activated carbon to eliminate other potential factors, such as lipids and charged molecules. This method of charcoal-stripping does change the neuronal response to glial shear-conditioned media, showing no difference from the static equivalent and a significant increase compared to the untreated shear condition (**Figure 4C, D**). Therefore, the responsible factor is likely either lipidic or a charged small molecule, typically positively charged, that would bind to the activated carbon.

**Figure 4.**
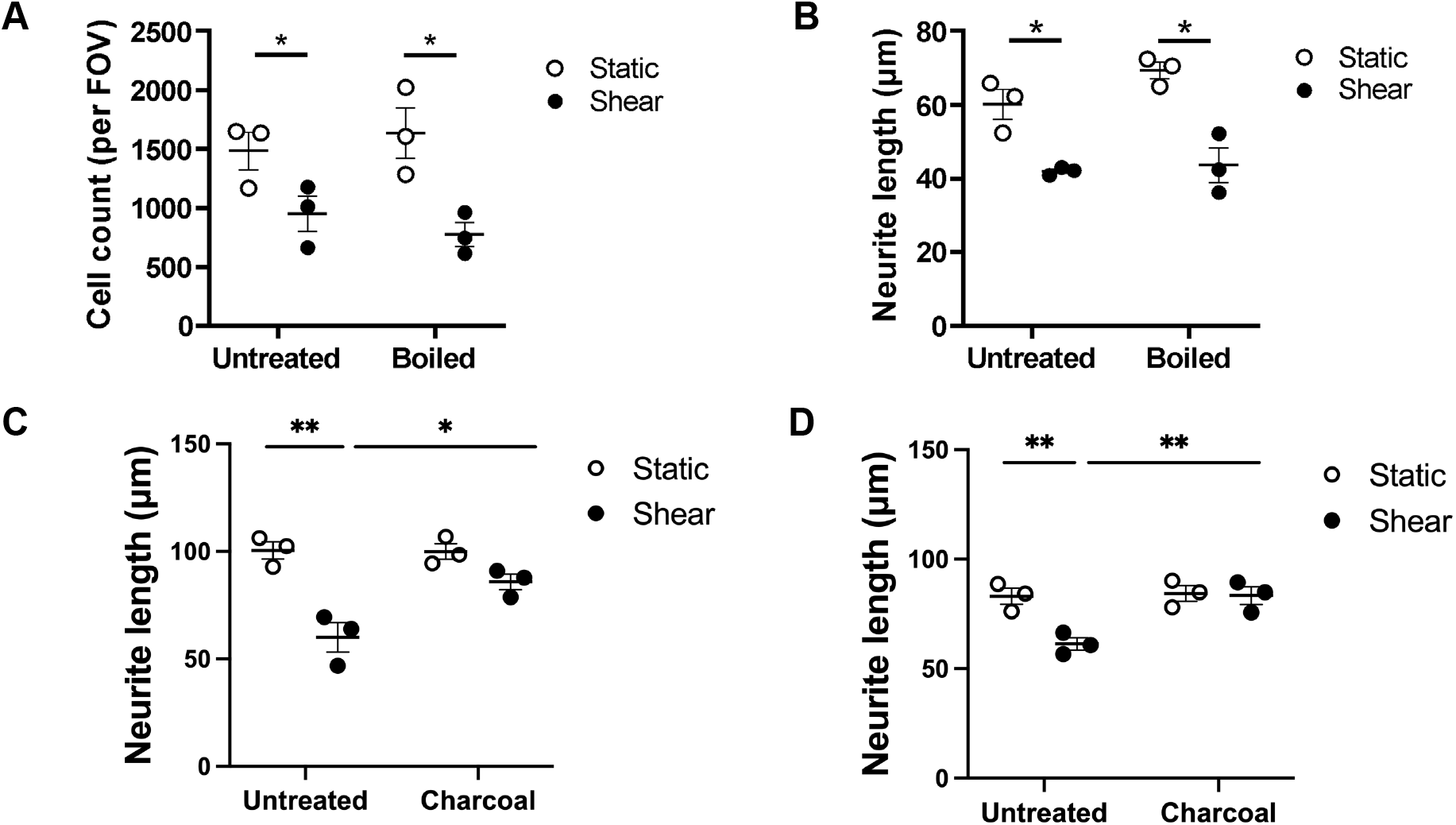
The toxic effect of glia under high shear is removed by charcoal stripping but not boiling. A) Quantification showing boiling the media from glial co-cultures has no effect on the shear-induced reduction in neuronal cell count. B) Quantification showing boiling co-culture media also has no effect on shear-induced neurite reduction. C-D) Quantification showing charcoal-stripping alleviates effect of shear-induced neurite reduction in neurons treated with astrocyte (C) and microglia (D) conditioned media. Groups for static and shear were compared by ratio paired t-tests. *p<0.05, **p<0.01.

### Shear-induced increase in glial eATP release is responsible for neurotoxicity effects

We next sought to identify a soluble factor that could feasibly be released by both astrocytes and microglia under high shear that promotes neuronal toxicity. Previous studies had shown the sphingolipid known as sphingosine-1-phosphate (S1P) promotes neuronal retraction via interaction with sphingosine–phosphate receptor 3 (S1PR3), and this also mediates inflammatory responses in astrocytes (38,39). We therefore first investigated if S1PR3 on neurons plays a role in neurite inhibition/retraction in response to glial shear-conditioned media. Pharmacological inhibition of S1PR3 in neurons prevented the reduction in neurite outgrowth for microglia-conditioned media but only showed modest, non-significant effects with astrocyte-and mixed glial-conditioned media (**Supplemental Figure S2A**). We were able to recreate prior experiments showing that treatment of neurons with 1 μM S1P does cause significantly shorter neurites compared to a vehicle control (**Supplemental Figure S2B**) (38). Nonetheless, inhibiting sphingosine kinase in glial cells did not prevent the shear-induced neurotoxicity (**Supplemental Figure S2C**), and there was no significant difference in S1P levels in shear-conditioned media compared to static control (**Supplemental Figure S2D, E**). Given these findings, we investigated other molecules that may be adsorbed by charcoal.

Because we observed an elevated release of eATP from both glial cell types under fluid shear stress, we next tested if charcoal filtration removed the negative charged eATP from the media. Charcoal filtration removes nearly all eATP from both astrocyte- and microglia-conditioned media (**Figure 5A, B**). Given this surprising result, we investigated whether ATP is the soluble factor responsible for the neurite inhibition/retraction. eATP binds to numerous purinergic receptors, including the metabotropic P2Y receptor family and the ionotropic P2X receptor family. We focused on P2×7R given its established role in neurological disorders (40–42). After confirming that differentiated SH-SY5Y cells express P2×7R (**Supplemental Figure S3**), we pharmacologically inhibited the P2×7 receptor in neurons while treating with the astrocyte- and microglia-conditioned media, as per previous experiments. Blocking P2×7R effectively prevents the effect of neurite reduction observed with glial shear-conditioned media (**Figure 5C, D**), demonstrating that the neuronal toxicity effect is mediated by increased extracellular ATP binding to the P2×7 receptor on neurons (**Figure 5D**). Collectively, our data show pathological fluid shear stress provokes an elevated release of extracellular ATP from both astrocytes and microglia, which can activate the low-affinity P2×7 receptor in neurons to cause downstream neurotoxic effects (**Figure 5E**).

**Figure 5.**
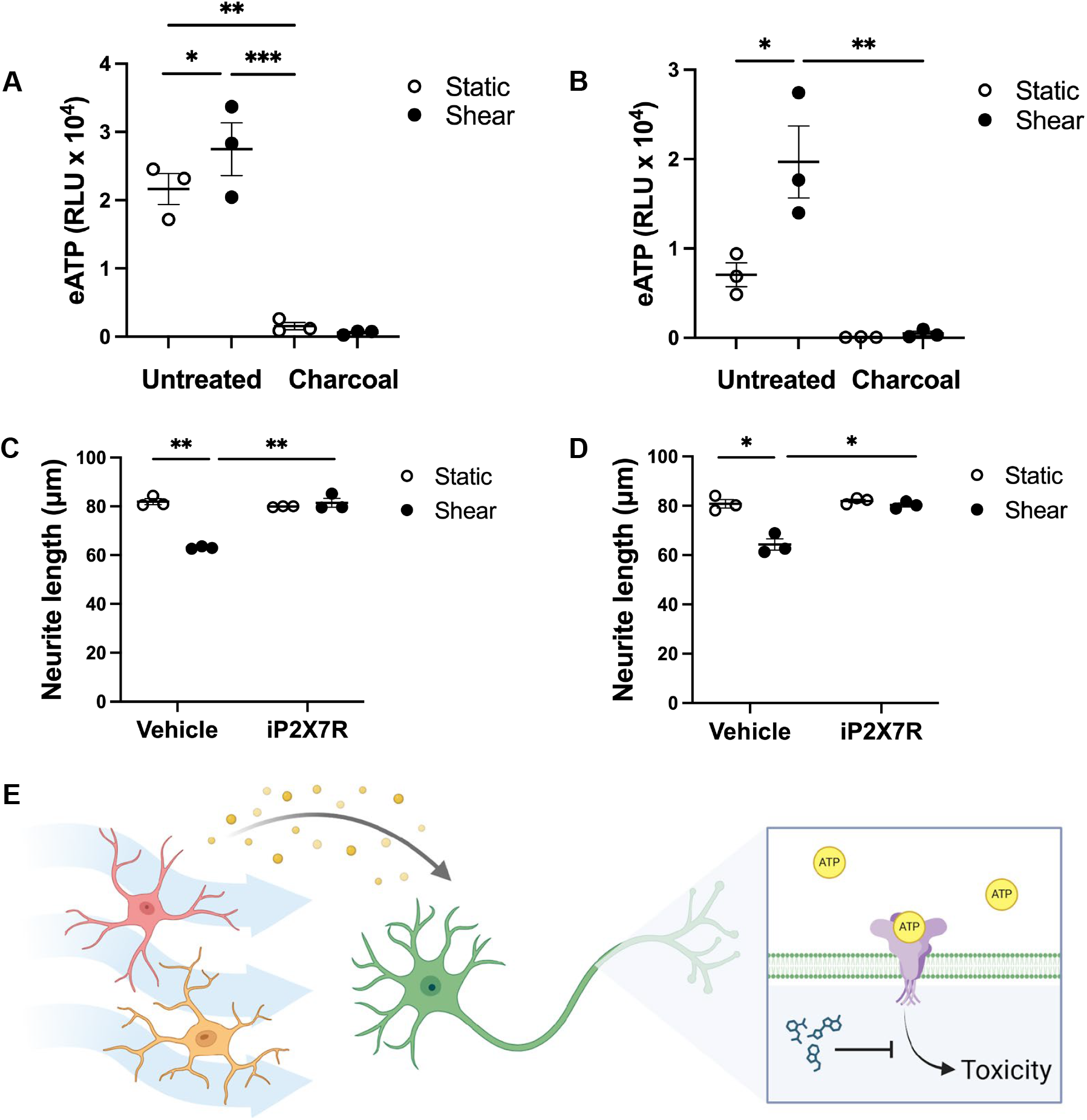
Shear-induced increases in glial eATP release is responsible for neuronal toxicity effects. A-B) Charcoal stripping removes all eATP from the astrocyte (A) and microglia (B) shear-conditioned medium, presented in relative luminescence units (RLU). C-D) Quantification showing that blocking purinergic receptor P2×7 in neurons abrogates the neurite reduction effect of astrocyte (C) and microglia (D) shear-conditioned media. Groups for static and shear were compared by ratio paired t-tests. *p<0.05, **p<0.01. E) Graphic summary of how high fluid shear stress on astrocytes and microglia stimulates ATP release, which signals to neurons to induce toxicity. This effect can be inhibited by blocking the purinergic receptor P2×7R on neurons. Made with an institutional license for Biorender.com.

## Discussion

Biophysical fluid forces are a pervasive component of tissue physiology, including for cells outside of blood vessels. Investigating how neural cells sense and respond to fluid forces will advance the understanding and treatment of pathologies associated with altered fluid flow, such as traumatic injury, stroke, and glioma. We show that astrocytes and microglia exhibit some differential responses to pathological fluid shear stress, with astrocytes displaying increased metabolic activity and glutamate release, while microglia show increased accumulation of lipid droplets. Important to our work, both astrocytes and microglia release significantly more extracellular ATP under pathological levels of fluid shear, which we show promotes downstream effects on neuronal cell survival and neurite outgrowth through purinergic P2×7R signaling.

Glial cells have been known to respond to mechanical cues like stiffness and stretch, which alters their functionality and phenotype in the context of injury and electrode implantation (43,44). Recent studies have also started to evaluate how glial cells respond to fluid shear stress. At physiological levels, astrocytes were shown to sense fluid shear forces relevant to the glymphatic system through the mechanosensitive receptor Piezo1 (13). At more pathological levels, both astrocytes and microglia were shown to adopt pro-inflammatory changes in response to high fluid shear stress *in vitro* (45), which may also have functional implication *in vivo* (46). Our findings further validate that glial cell functionality is altered by high fluid shear stress, showing it alters glial metabolism, metabolite secretion, lipid metabolism, and phagocytic activity in a cell-type specific manner. Given the established role of extracellular ATP as a danger signal (47,48), the convergent response of increased extracellular ATP release suggests that high fluid shear promotes cellular stress in both glial cell types. This response likely has widespread implications on the function of glial cells themselves but also other cell types in the brain such as neurons (47,48).

Our work starts to examine how sheared glial cells influence neuronal cell function, by treating differentiated SH-SY5Y neuron-like cells with glial shear-conditioned media. Our data show that glial cells under pathological shear release a soluble factor that significantly reduces neuronal cell survival and neurite outgrowth compared to static control glial cells. Ultimately, we show the soluble factor is extracellular ATP released by sheared glial cells, and the neurotoxicity effects are mediated by ATP binding to the receptor P2×7R on neurons, since blocking this receptor abrogated the effects. While there has been controversy in the field as to whether or not neurons express P2×7R *in vivo* (49), recent studies have validated the expression of this receptor and its relevance in neuronal functionality (50–52). Furthermore, while a prior study reported that SH-SY5Y cells downregulate P2×7R during neuronal differentiation (53), we validated protein expression in our cells by immunocytochemistry and others have confirmed receptor functionality in these cells (54). Differentiated SH-SY5Y neurons are commonly utilized for modeling neurons *in vitro*, but it should be noted that neuronal responses of this model may not accurately represent neuronal function *in vivo*, and future studies utilizing primary neuron cultures and *in vivo* models are necessary to fully characterize the effect of shear-induced glial-mediated neurotoxicity and the role of eATP release in this signaling cascade.

The P2×7 receptor has a lower affinity to ATP than other P2X family members and is only activated by relatively high levels of eATP (40–42). ATP concentrations in healthy tissue have been reported to be in the nanomolar range, whereas sites of inflammation can have ATP concentrations in the hundreds of millimolar range, well past the activation threshold of P2×7R in the low millimolar range (41). High ATP concentration or exposure for prolonged periods of time can lead to cytotoxicity and apoptosis, as we observed in neurons here. Because we observed this effect only after three days in shear-conditioned media, additional studies are needed to determine the short-term effects of sheared glial on neuronal cell function. In addition, while there was a significant increase in eATP between static- and shear-conditioned media for both glial cell types, the eATP levels of microglia at rest were nearly as high as that of astrocytes under fluid shear. It is likely, then, that other factors contribute to the neurotoxic effect, such as co-stimulation of other pathways like S1P binding to S1PR3. The ATP assay we used did not provide absolute concentration, nor would a concentration from this 2D fluid shear model provide definitive biological insight into the exact processes occurring within complex 3D tissue. Future studies should therefore evaluate the role of shear-induced signaling *in vivo* to better define and understand the tissue-level consequences. A further limitation lies in the use of immortalized cell lines, namely SH-SY5Y and microglia. These cell lines provide a useful model, but they neither fully recapitulate cellular responses of primary neurons or microglia *in vitro* nor replicate their behavior in an *in vivo* environment.

Traumatic injury, ischemic stroke, and glioma all involve disruptions to fluid flow in the central nervous system, which exposes cells to pathologically high fluid shear stress. We show here that even modest increases in fluid shear stress from physiological levels results in increased release of extracellular ATP by astrocytes and microglia, which promotes neurotoxicity via P2×7 receptor signaling in neurons. P2×7 receptor inhibition completely prevented the deleterious effects of high fluid shear stress, highlighting the role of this pathway in shear-induced neurotoxicity. Blocking P2×7 receptor signaling has been shown to improve outcomes in traumatic brain injury (25–27), preserve blood brain barrier integrity and alleviate pain post-stroke (28,29), and decrease glioma cell proliferation and tumor growth (30,31). Our results suggest that altered fluid dynamics could be a driver of purinergic signaling in these pathologies. These findings highlight the under-characterized role of mechanotransduction in glial-neuronal interactions, expands the current mechanistic understanding of neuropathologies associated with disrupted interstitial fluid dynamics, and provides further rationale for the use of therapeutic P2×7 receptor inhibition in the treatment of neuropathologies.

## Materials and Methods

### Cell Culture

Immortalized human microglia (ABM #T0251) were cultured up to passage p10 in Dulbecco’s Modified Eagle Medium (DMEM) supplemented with 10% fetal bovine serum (FBS) at 37°C and 5% carbon dioxide. Human cortical astrocytes (Sciencell #1800) were cultured up to passage p10 in Astrocyte Medium (AM) supplemented with 2% FBS and 1% astrocyte growth supplement. Cells were passaged every 7 days or at 80% confluency onto tissue culture-treated plasticware coated in 7.5μg/cm^2^ rat tail type I collagen. Media was changed every 2 days. SH-SY5Y neuroblastoma cells (ATCC CRL-2266™) were cultured in media containing 44.5% DMEM, 44.5% F12, 10% FBS, and 1% Non-Essential Amino Acids (NEAA) at 37°C and 5% carbon dioxide. Cells were passaged at 80% confluency. Media was refreshed every 2 days. Differentiation into neuron-like cells was initiated 24 to 48 hours after plating cells. Media was replaced with Neurobasal™ medium supplemented with 0.25% GlutaMAX, 1% B-27™ Supplement, 0.5% N-2 Supplement, and 10 μM all-trans retinoic acid (ATRA). Neurite outgrowth was monitored for 5 days, refreshing the media every 48 hours.

### Application of Fluid Shear Stress

Microglia and astrocytes were plated in 6-well tissue culture-treated plates coated in 7.5μg/cm^2^ rat tail type I collagen at a density of 1×10^5^ cells per well. 24 hours after plating, the cells were serum starved for three hours in basal Astrocyte Media (AM) supplemented with 1% B-27™ Supplement and 0.5% N-2 Supplement. Experimental cells were exposed to laminar shear stress on an orbital shaker at 100 rpm, equivalent to a low pathological shear stress of 0.1 dynes/cm^2^, for 18 hours in a cell incubator at 37°C and 5% carbon dioxide. Control cells remained static in the same incubator. After exposure to shear stress, conditioned media was collected and stored at -80°C, and cells were washed in phosphate-buffered saline (PBS) and fixed at room temperature with 4% paraformaldehyde for 20 minutes.

### Live/Dead Assay

Human astrocytes, microglia, or both were plated in collagen-coated 6-well plates at 50,000 cells each per well, allowed to adhere overnight in complete astrocyte medium, and serum starved in basal astrocyte medium containing 1% B-27™ and 0.5% N-2 supplements for three hours prior to the start of shear. Shear culture was conducted as described above for 18 hours, with a separate plate kept in static conditions as a control. The cells were labeled with NucRed Live 647 (Thermo Fisher) according to the manufacturer’s protocol, which labeled all cells red. The dead cells were labeled using 4’,6-diamidino-2-phenylindole (DAPI) diluted 1:5,000 in 1X PBS for 5 minutes. The samples were then imaged at three to five locations around the periphery of each well (maximum fluid shear stress), and the images were analyzed using ImageJ software. Both the total number of cells and percentage of dead cells to live cells were quantified.

### Metabolic Assay

Cellular metabolic activity was measured using alamarBlue HS (Thermo) according to the manufacturer’s instructions. Briefly, after 18 hours of culture under shear, the condition media were replaced with alamarBlue HS diluted in serum-free media and incubated for 4 hours. Media fluorescence was measured using a PerkinElmer Enspire plate reader, with autoclaved alamarBlue reagent serving as a positive control. Separately, the cells were lysed using 300 μL of RIPA Lysis and Extraction Buffer (Thermo) per each well of a 6-well plate that was distributed using a cell scraper. The DNA content per well was then quantified using the Quant-iT™ PicoGreen™ dsDNA Kit (Thermo), performed according to manufacturer’s instructions. Sample fluorescence was measured using a PerkinElmer Enspire plate reader using excitation of 480 nm and emission of 520 nm. Data from the alamarBlue assay (in relative fluorescence units) was then normalized to total nanograms of DNA content per well.

### Metabolite Quantification

Separately, shear-conditioned media were used for metabolite quantification. Extracellular ATP was measured via RealTime-Glo™ Extracellular ATP Assay (Promega GA5010) according to the manufacturer’s specifications. Glutamate and lactate released by astrocytes and microglia were detected in conditioned media via Glutamate-Glo™ Assay (Promega) and Lactate-Glo™ Assay (Promega), also performed according to manufacturer’s specifications. Briefly, glial conditioned cell culture media was collected from overnight cultures. Media were used undiluted for the ATP assay; samples were diluted 1:10 for the glutamate assay and 1:100 for the lactate assay. The media were mixed with the detection reagent at a 1:1 ratio in a 96-well plate with opaque sides, incubated for 1 hour at room temperature, and luminescence was read with a PerkinElmer Enspire plate reader.

### Phagocytic Activity Assay

Microglia were cultured and treated with fluid shear stress under the same conditions as above. After 18 hours of exposure to 0.1 dynes/cm^2^ fluid shear stress, cells were incubated with 2 μL of FluoSpheres™ 1 μm Polystyrene Microspheres (Thermofisher) suspended in 2 mL of DMEM supplemented with 10% FBS for 6 hours under static conditions, after which the microsphere-containing media was removed and cells were washed three times with 1X PBS to remove any excess microspheres. Cells were stained for 15 minutes in a cell incubator with 0.2 μL CellTracker™ Red (Thermofisher) fluorescent probe in the same media as before. After 15 minutes, the media were replaced with fluorescent probe-free media and incubated for 15 additional minutes. Cells were washed with 1X PBS and fixed with 4% paraformaldehyde for 20 minutes. Cells were then labelled with 4’,6-Diamidino-2-Phenylindole (DAPI) (1:5000, Invitrogen) for 5 minutes. DAPI was removed, cells were washed with PBS, and the samples were stored at 4°C until imaging. Images were acquired at 20X using a Keyence BZ-X800 fluorescence microscope.

### Lipid Droplet Staining

Lipid droplets were stained in live cells prior to fixation using BODIPY 493/503 dye (Invitrogen). The channel of an Ibidi µ-Slide I Luer microfluidic chip was coated with 1 mg/mL rat tail collagen for 30 minutes at 37°C, rinsed with 1X PBS, and loaded with astrocytes or microglia at 300,000 cells/mL. After allowing for cell attachment overnight, the device was connected to a Chemyx Fusion 200 microsyringe pump using polytetrafluoroethylene tubing (ID 0.012, thickness 0.009, Weico). Serum-free Astrocyte Media (AM) supplemented with 1% B-27™ Supplement and 0.5% N-2 Supplement was pumped through the microfluidic chip at a rate of 0.08 mL/minute, equating to a shear stress rate of 0.1 dyne/cm^2^, for one hour. After shear, cells were stained with BODIPY 493/503 dye 1 mg/mL in DMSO stock solution diluted 1:5000 in cell culture media for 30 minutes at 37°C, rinsed with 1X PBS, and fixed with 4% paraformaldehyde for 20 minutes. Images were acquired at 20X using a Keyence BZ-X800 fluorescence microscope.

### Treatment of Neurons with Conditioned Media

Neuroblastoma cells were plated in 48-well tissue culture-treated plates at a density of 7.5×10^3^ cells per well for differentiation, as described above. After five days of differentiation, media was replaced with 200 μL of glial conditioned media (static or shear, from astrocytes or microglia) in each well. To denature proteins, some conditioned media were boiled at 95°C for 5 minutes in a water bath and cooled on ice before use. Alternatively, conditioned media samples were filtered by adding 1 mg/mL of activated charcoal (−4+8 mesh, Thermo Fisher) and mixing for two hours on a tube rotator at 10 rpm. Charcoal filtered media was then passed through a 0.22 μm filter prior to use. Neurons were incubated with conditioned media for three days, after which the media was aspirated, and cells were fixed with 4% paraformaldehyde for 20 minutes, and stored in PBS at 4°C.

### Immunocytochemistry

Fixed neurons were washed three times with PBS (ten minutes each), then incubated in a blocking buffer containing 0.3% Triton X100 and 3% Normal Donkey Serum in 1X PBS for one hour at room temperature. Primary antibody for beta-III Tubulin (1:1000, Abcam #ab7751) was diluted in blocking buffer, and cells were incubated with primary antibody overnight at 4°C. The following day, cells were washed three times with 1X PBS (ten minutes each) at room temperature, then labeled with a secondary antibody (1:500) in blocking buffer for one hour. The antibody solution was then aspirated, and cells were washed three times with PBS (10 minutes each), followed by counterstaining with DAPI (1:5000, Invitrogen) for 5 minutes. DAPI was removed, cells were washed with 1X PBS, and the samples were stored at 4°C until imaging. Images were acquired at 20X using an EVOS 6500-FL fluorescence microscope (Thermofisher).

### Image Analysis

All images were analyzed using ImageJ (National Institutes of Health). Microglia phagocytic activity was quantified by determining the total number of cells per field of view and quantifying the number of cells that took up microspheres, comparing between fluid shear-exposed cells and static cells. Lipid droplets were quantified by measuring corrected total cell fluorescence and comparing the integrated density between shear-treated and static cells. Neuronal cell counts were performed using the Cell Count plugin on the DAPI image. Neurite length was quantified using the plugin NeuronJ, tracing an average of 30 axons per image.

### Statistical Analysis

All data were analyzed and graphed using GraphPad Prism software (GraphPad v.10.2.2). All experiments included n=3, unless otherwise specified. Neurite length and cell count data were analyzed via ratio-paired two-tailed t-tests. For all graphs, error bars represent the standard error of the mean (SEM), and each data point represents a biological cell culture replicate. Statistical significance is indicated by *p < 0.05, **p < 0.01, ***p < 0.001, ****p < 0.0001.

## Conclusions

Glial cells play diverse roles both in maintaining neural homeostasis and in the development and resolution of neuropathology (1,4,6). These cells reside in the interstitial space of the brain parenchyma, which is subjected to continuous flow of interstitial fluid (9). Disruptions to this fluid flow are associated with a multitude of neuropathologies (17–19). In this study we identify that high fluid shear stress modulates glial cell functionality with neurotoxic consequences. Using an *in vitro* 2D model of fluid shear stress, we demonstrate that fluid shear stress rates of 0.1 dynes/cm^2^ results in functional changes in both astrocytes and microglia, with distinct responses between cell types. High shear stress promotes an increase in the release of extracellular ATP in both astrocytes and microglia, which in turn promotes neuronal toxicity via P2×7 receptor signaling. Further research will elucidate the role of high fluid shear stress in neurotoxicity, including the mechanism downstream of P2×7 receptor signaling in neuronal cells. The results of this study highlight an under-characterized role of mechanobiological processes in glial-neuronal communication and provide further support for the therapeutic efficacy of P2×7 receptor inhibition in neurological disease.

## Acknowledgements

The authors would like to thank Jim Chambers in the UMass Nikon Center of Excellence Light Microscopy Facility for assistance on training and imaging guidance throughout the project. We would also like to acknowledge Aleeza Zilberman for assisting with data collection for the live/dead experiments.

## Authorship Contributions

RCC conceptualized the project idea; ST, CS, and RCC developed the investigational plan and methodology; ST, CS, ZK, and AB conducted experiments for data acquisition and analysis; RCC supervised experimental design and data analysis; ST and RCC developed the original manuscript draft and curated data for publication; RCC acquired funding and managed the procurement of reagents and supplies; all authors edited and reviewed the final manuscript.

## Funding

This work was supported in full by start-up funds provided by the University of Massachusetts Amherst.

## Data Sharing and Data Availability

The data supporting the presenting findings are publicly available at Open Science Framework and can be provided upon request to the corresponding author.

## Declaration of interests

The authors declare no conflicts of interest.

## Ethics Statement

The present studies did not involve the direct use of animals or human subjects and therefore were not subject to ethical regulatory guidelines.

## Notes

### Competing Interest Statement

The authors have declared no competing interest.

