## Supplemental Figures for "Fluid shear stress on astrocytes and microglia promotes neuronal toxicity *in vitro* via purinergic signaling"

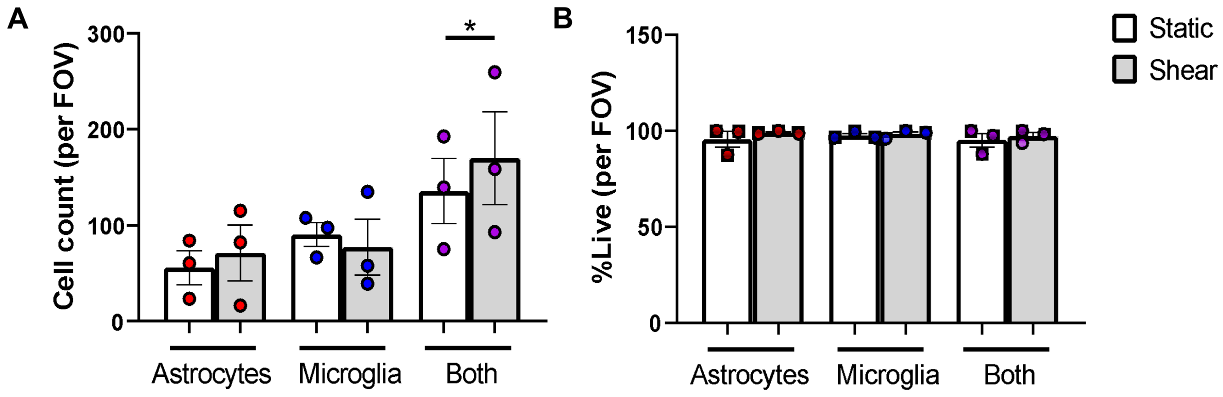


**Figure S1: Shear stress of 0.1 dynes/cm$^2$ does not affect glial cell survival.** A) Quantification of average cell counts after overnight culture under static and shear conditions, based on DAPI staining. B) Quantification of live cells as a percentage of total cells, based on NucRed Live and DAPI staining. *$p<0.05$.

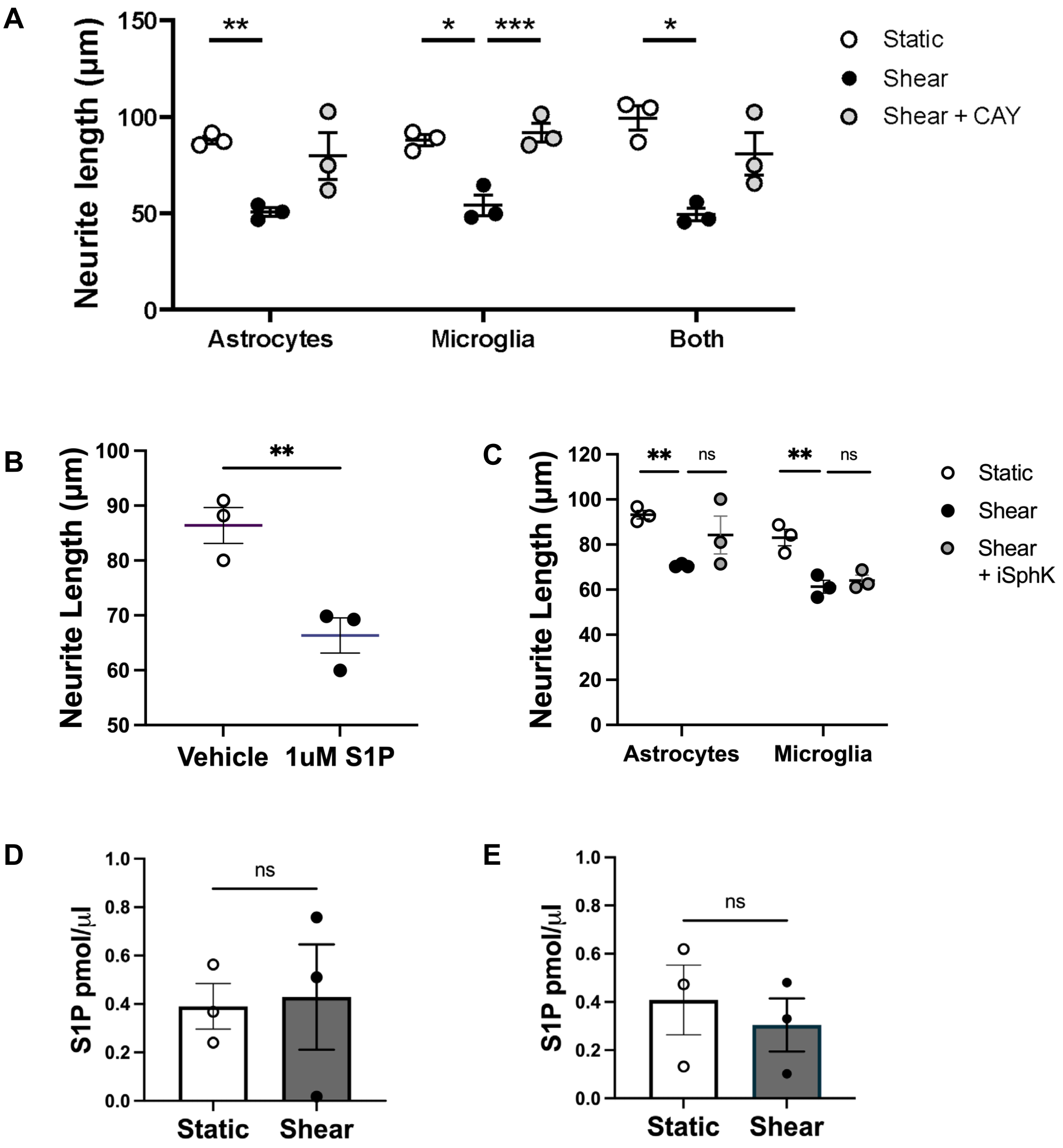


**Figure S2: Effects of blocking S1P-S1PR3 pathway on glial shear-induced neuronal toxicity.** A) Quantification showing that blocking S1PR3 in neurons prevented the effect of shortened neurite length in neurons treated with microglia shear-conditioned media. There was a similar but not statistically significant effect with astrocyte or glial co-culture conditioned media. B) Quantification showing the addition of S1P to neuronal cell culture media resulted in shortened neurites. C) Quantification showing that blocking sphingosine kinase 1 (SphK) in glial cells did not prevent the effects of shear-conditioned media. D-E) Quantification of S1P in astrocyte (D) and microglia (E) conditioned media, which does not change with fluid shear. Groups for static and shear were compared by ratio paired t-tests. *$p<0.05$, **$p<0.01$, ***$p<0.001$.

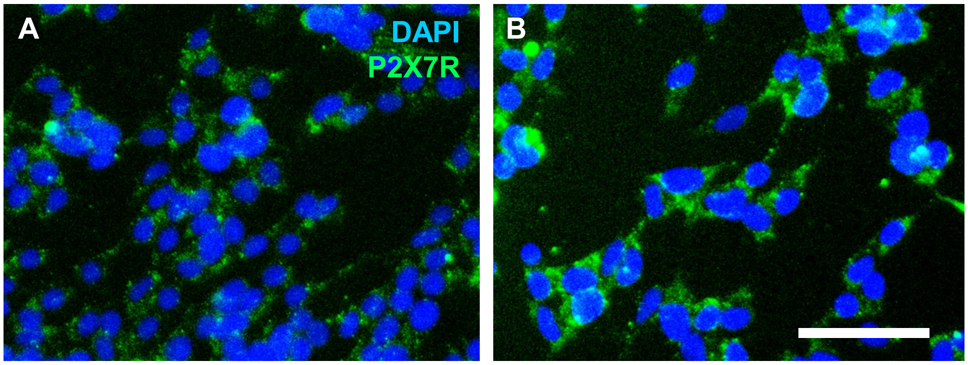


**Figure S3: Expression of P2X7R in SH-SY5Y before and after neuronal differentiation.** A) Representative immunofluorescence image of undifferentiated SH-SY5Y cells labeled for P2X7R. B) Representative immunofluorescence image of SH-SY5Y cells after 5 days of neuronal differentiation labeled for P2X7R. Scale bar is 50 µm.
